# Quantifying the demographic vulnerabilities of dry woodlands to climate and competition using range-wide monitoring data

**DOI:** 10.1101/2020.04.03.024497

**Authors:** Robert K. Shriver, Charles B. Yackulic, David M. Bell, John B. Bradford

## Abstract

Climate change is expected to alter the distribution and abundance of tree species, impacting ecosystem structure and function. Yet, anticipating where this will occur is often hampered by a lack of understanding of how demographic rates, most notably recruitment, vary in response to climate and competition across a species range. Using large-scale monitoring data on two dry woodland tree species (*Pinus edulis* and *Juniperus osteosperma*), we develop an approach to infer recruitment, survival, and growth of both species across their range. In doing so, we account for ecological and statistical dependencies inherent in large-scale monitoring data. We find that warming and drying conditions generally lead to declines in recruitment and survival, but there were some idiosyncrasy in the strength of responses across species. Climate conditions lead to vulnerable regions, such as *Pinus edulis* in N. Arizona, where both survival and recruitment are low. Our approach provides a path forward for leveraging emerging large-scale monitoring and remotely sensed data to anticipate the impacts of global change on species distributions.

## Introduction

Changing climate, disturbance regimes, and human activity are expected to reshape the distribution of forest and woodland species across the globe, potentially transforming the structure of these ecosystems (Allen et al. 2010, McDowell et al. 2018). Although, mortality of overstory plants are often the most obvious indicators of declining forest and woodland health (e.g. Millar and Stephenson 2015), the resilience and long-term viability of tree species in the face of environmental change will be determined by not only the survival of existing individuals, but also the recruitment and growth of new individuals that drive population recovery and spread, i.e. resilience (Jackson et al. 2009). Evidence of declining forest health and resilience due to declining recruitment is increasingly common (Petrie et al. 2017, Stevens-Rumann et al. 2018). Yet anticipating where forest and woodland species may be most vulnerable to environmental change is often hampered by a lack of understanding of how rates of survival, growth, and, most notably, recruitment vary across a species range and the environmental conditions driving them.

Demographic processes are increasingly recognized to be critical to understanding species range shifts and ecosystem state changes in response to climate change (Briscoe et al. 2019). But, efforts to estimate how plant recruitment, survival, and growth vary across large spatial scales are limited, in part because many traditional demographic inference approaches (e.g. Easterling et al. 2000) do not typically accommodate diverse data structures and the ecological/statistical dependences inherent to large spatial data. Instead, plant demographic analyses have typically placed the onus on researchers to mark and return to individuals and independently measure each demographic transition in the field, making it logistically challenging to scale data collection to larger areas. At the same time, there has been an explosion of diverse datasets from large-scale field monitoring and remote sensing (e.g. Forest Inventory and Analysis Database, lidar) with the potential to revolutionize our understanding of spatio-temporal plant demographic processes and their population consequences. Yet, these data sources rarely provide data on both survival and recruitment at the individual-scale resolution required for traditional demographic modeling approaches. Harnessing the power of these datasets will require flexible modeling approaches that can link detailed, individual demographic data with additional data sources that describe how demographic processes drive changes in abundance and population structure across large landscapes and over decades (Shriver et al. 2019).

Here, we develop an approach to infer the range-wide recruitment, survival, and growth rates of two widespread dry woodland species using large-scale Forest Inventory and Analysis (FIA) data. Because FIA plots encompass nearly the entire range of many tree species, they present a unique opportunity to understand how demographic rates vary across a species’ range and identify where populations may be most vulnerable to changing climate and disturbance. But, FIA data also present a number of challenges (see Methods for full explanation), most notably accounting for ecological dependencies in quantifying recruitment. Specifically, seedlings are not tagged but simply counted, thus the fate (growth/survival) of existing seedlings are not independently quantified from new recruits. We overcome this challenge by developing an integrated population modeling approach that accounts for ecological dependencies while also accounting for spatial autocorrelation and sharing information across sites in a rigorous way.

## Methods

### FIA Data

We developed our demographic models using the publicly available FIA database (http://www.fia.fs.fed.us/). FIA is a systematic and standardized survey of forested regions in the entire United States, including both public and private lands. Full details on the sampling design can be found in Bechtold and Patterson (2005).

We focus our analyses on two widespread dry woodland species in the Colorado Plateau and Great Basin regions: *Pinus edulis* (hereafter PiEd) and *Juniperus osteosperma* (hereafter JuOs). Nearly the entire ranges of both species are within the United States, thus FIA data provide a near complete survey of their range-wide dynamics. Because our primary focus was quantifying demographic rates of each species and linking these to climate across their range, we excluded all plots in which fire mortality or tree harvesting occurred. This resulted in 2,013 plots with 16,951 tagged PiEd individuals, and 2,380 JuOs plots with 25,105 tagged JuOs individuals. All PiEd and JuOs individuals greater than 15.24 cm (6 in.) in height are surveyed. Within each plot, all adult trees greater than 12.7 cm (5 in.) diameter are assigned unique tags and tracked within 4, 7.32 m (24 ft.) radius subplots. All saplings <12.7 cm & > 2.54 cm (1 in.) diameter are assigned unique tags and tracked within 4, 2.07 m (6.8 ft.) radius microplots within the larger adult plots. Finally, seedlings <2.54 cm diameter are counted within the same microplots as the saplings.

Two censuses were conducted 10 years apart in each plot. In some cases, additional plot surveys occurred between the standard 10 year interval. These additional surveys were excluded because they were sporadic and not standardized across the dataset. The exact timing of the initial censuses varied by state and region within state (typically 10-20% of plots in each state are surveyed each year) but occurred between 2000 and 2007.

### Demographic Modeling

Data on adult and sapling growth and survival are collected at the individual level. Individual plants >2.54 cm diameter are tagged, and as a result the growth and survival of an individual plant can be tracked over the census interval. We develop growth (i.e. change in size) and survival models following the well-developed generalized linear model functional forms common for plant demography models (Rees et al. 2014). Individual diameter size change is modeled as

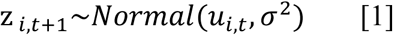

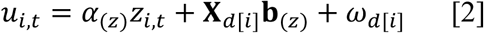

Where Z_*i,t*_ is the size of plant *i (i=1…I)* in the first census (*t*), α_(*Z*)_ is a regression coefficient for plant size, **X**_*d*[*i*]_ is a vector of plot-level environmental covariates for plot *d* (*d=1…D)* where individual *i* is located, **b**_(*Z*)_ is a vector of environmental regression parameters specific to the size model, ω_*d*[*i*]_ is a plot-level spatial random effect, and σ is a variance parameter.

Similarly, survival probability is modeled as

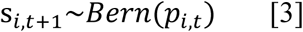

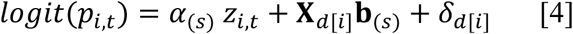

where *p*_*i,t*_ is the probability of survival for individual *i* from *t* to *t+1, z*_*t,i*_ is again the size of plant *i* in the first census (*t*), α_(*s*)_ is a regression coefficient for plant size on survival, **b**_(*s*)_ is a vector of environmental regression parameters specific survival, and δ_*d*[*i*]_ is a plot-specific spatial random effect for each individual *i*. Note, the comparatively small number of observed JuOs mortality events led spatial random effects to be non-identifiable, thus were not included in the JuOs survival model.

Spatial random effects were fit using a predictive process model (Banerjee et al. 2008, Latimer et al. 2009). Predictive process models address the computational challenges of fitting spatial models to large datasets by reducing point locations (i.e. plots) to a lesser number of constituent knots that encapsulate the landscape of spatially autocorrelated processes not explained by covariates. In the case of growth random effects,

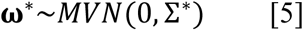

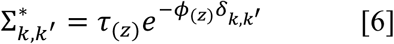

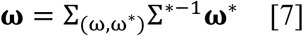

**ω**\* is a K-length vector of random effects (**ω**\***=**ω*_1_, ω*_2_…ω*_*k*_) associated with each knot (*k*). ∑* is a covariance matrix where each element is a correlation among knots weighted by distance, δ*k,k*^′^. *ϕ* is a parameter describing the rate at which correlations decay as a function of distance (km), and τ is an error term. **ω** is a D-length vector of random effects for each plot (ω_*d*_ = ω_1_, ω_2_ … ω_*D*_). The underlying knot-based spatial landscape is then linked back to specific plots using Eq. 7, where ∑_(ω,ω_^*^) is a cross-covariance matrix which describes the spatial relationship between plots (**ω**) and knots (**ω**\*) using Eq. 6, where in this case δ*k,k*^*′*^ is the distance (km) between each the fuzzed location of plot (*d*) and knot (*k*) pair. While model fit will improve as the number of knots increase, the choice of the number of knots is a tradeoff of model fit and computational efficiency. We follow the recommendations of Latimer et al. 2009 (i.e. 100-400 knots) by using 200 knots who’s locations are assigned to maximize coverage of FIA plot locations using the “cover.design” function in the “fields” package (v. 9.6) in R (Nychka and Furrer 2017).

FIA data present several challenges for estimating recruitment rates. First, unlike the growth and survival of saplings and adults, the recruitment, growth, & survival of seedling is never directly observed. All conifers <2.54 cm diameter but >15.25cm height are simply counted, making it impossible to directly separate new recruits from the fate of pre-existing seedlings. In other words, the change in the count from census to census represents both new recruitment, but also the survival and growth of existing plants. While this data structure does not preclude inference on the underlying reproduction rate, it is incompatible with most traditional statistical approaches (e.g. Poisson GLMs) for estimating plant reproduction recruitment, which require clearly identifying the reproductive output (e.g. seeds) of existing individuals and the fate of these propagules.

Second, the search area for seedlings is small given the low density of seedlings, and considerably smaller than area adult trees are measured over (∼12x smaller). This presents a challenge for estimating recruitment because as the number of individuals in a plot declines separating the true signal of environmental and spatial processes from noise introduced by sampling and demographic stochasticity is increasingly difficult. This may be particularly problematic for tree species exhibiting spatially and temporally heterogeneous seedling distributions, such as woodland tree species (Bell et al. 2014).

To address these challenges, we developed an integrated size-structured population modeling approach that shares available information among our different datasets (i.e. growth/survival of adults/saplings and counts of seedlings) and across FIA plots to infer the growth and survival of seedlings along with the reproductive output of existing trees leading to new recruits. Because the fate of all tagged individuals in the first census (i.e. any individual >2.54 cm) is already known, our goal is to build a model that describes the fate of all untagged individuals. Untagged individuals include all plants <2.54 cm and any plants that were not tagged in the first census, but reached the minimum tagged size (2.54 cm) by the second census. Plants reaching the 2.54 cm threshold in the second census could comprise existing plants previously <2.54 cm or new recruits.

We model the number of untagged plants in a plot as conditionally Negative Binomial

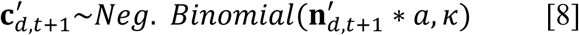

Where 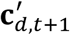 is a 5×1 vector of the counts of all untagged plants in plot *d* at the second census *t*+1. 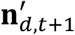 is a 5×1 vector of area standardized occurrence rates. *a* is the total plot area in which plants were counted. And, *κ* is a dispersion parameter. Each element in 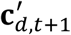 represents the counts of individuals in each 2.54 cm (or 1 inch) diameter class up to 12.7 cm. While the choice of diameter class is flexible, we use the natural choice of 2.54 cm diameter classes for all trees because the FIA dataset already lumps any plants <2.54 cm into a single class. Only plants less than 12.7 cm (i.e. size classes 1 to 5) were considered because untagged plants larger than this are more likely a results of previous missed observations than growth and recruitment from the smallest classes. 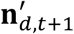 is defined as

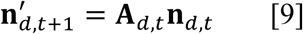

Where **A**_*d,t*_ is a 5 × 30 (h=1…5, j=1…30) discretized integral projection model (IPM) kernel (i.e. a matrix projection model with 2.54 cm size classes) describing all the pathways by which an existing plant could lead to an untagged plant (survival/growth of existing plants <2.54 cm and new reproduction from existing plants). **n**_*d,t*_ is a 30×1 vector of area standardized rates of occurrence of all plants in the first census in each 2.54 cm bin, derived from the empirical counts of all plants.

**A**_*d,t*_ is made up by the two different pathways by which untagged plants may appear: 1) the growth/survival of plants that were <2.54 cm in the previous census, and 2) recruitment arising from reproduction of existing trees.

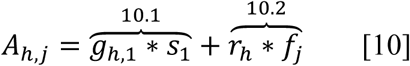

The first term (10.1) describes the fate of individuals <2.54 cm at time *t. g*_*h*,1_ aregrowth transition probabilities describing the movement of individuals in size class 1 to size classes 1 to 5. *s*_1_ is the survival probability of individuals in size class *j*=1 at time *t*. The second term (10.2) describes the number of new recruits produced per existing plant in each size class, *f*_*j*_, and the probability new recruits will transition to size class *h* by the second census. Note each element in Eq. 10 would also be indexed by site (*d*) and census interval (*t*), but we have omitted this for clarity.

Like tagged plants, *g*_*h*,1_ is defined by a normal distribution, but here it is a discretized kernel to account for size class binning. For a given site, *d*,

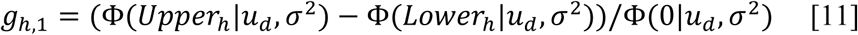

Where Φ indicates a cumulative normal distribution with mean *u* and variance σ^2^ evaluated at the upper and lower size limit of size bin *h* (Doak et al, In Revison). The final term renormalizes the kernel to strictly positive size values to prevent biologically impossible transitions to negative sizes. *u* is the same function used to evaluate the growth of tagged individuals (see eqs. 1 and 2), but in this case individual size (*z*_*i,t*_) is approximated by the midpoint of bin *j=1, m*_1_

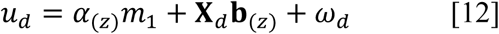

*s*_1_ is also based on the same survival function used for tagged plants, again approximated by the midpoint of bin *j=1* (*m*_*j*_).

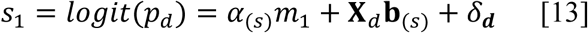

*r*_*h*_ uses a Gaussian kernel (normalized to sum to 1) to estimate the probability of any new recruit reaching size classes 1 to 5 as

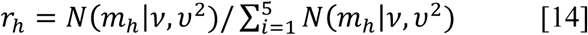

Finally, the function of primary interest is the number of new recruits produced per existing tree of size j, *f*_*j*_.

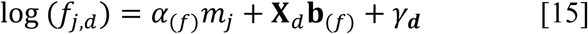

Where *f*_*j,d*_ is the number of new recruits produced per adult of size *j* at across all 30 size classes. *m*_*j*_ is the midpoint of each of the size classes, and α_(*f*)_ is a regression parameter describing the effect of size on reproduction. **b**_(*f*)_ are regression parameters describing the impact of size on reproductive output, and γ_*d*_ is a spatial random effect again defined by a predictive process (see Eq. 5) with its own parameters *ϕ*_(*f*)_ and τ_(*f*)_.

### Covariates

Based on previous research in dry woodland ecosystems, covariates in **X**_*d*_ were selected to summarize the effects of moisture availability (MA), heat stress (HS), and neighbor density (ND) on the growth, survival, and recruitment (McDowell et al. 2008, Allen et al. 2010). Moisture availability was the mean growing season (May to October) available soil water (i.e. >-3.9 MPA) over 40 to 100 cm depth over the 10-year census interval. Heat stress was the average temperature over the growing season over the 10-year census interval. Neighbor density was the basal area density of all living trees in the plot at the first census.

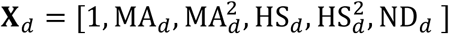

Squared MA and HS terms were added to account for possibility of nonlinear responses of species to environmental conditions across their range. Although multi-model inference using differing variables and functional form is possible given unlimited time (models take about 5-8 days to fit), we chose instead to focus on a limited set of variables and functional forms that are common to the demographic modeling literature and well supported based on our prior knowledge of the biology of Piñon and Juniper. All covariate parameters were given non-informative priors. Further details and covariates, model fitting, and priors can be found in the Supplemental Material.

## Results

To understand how vital rates vary across climate and geographic space we estimated posterior mean demographic rates for a 15 cm diameter individual using the observed moisture availability, heat stress, and neighbor density in each plot as well as plot random effects. This allowed us to quantify and visualize how vital rates change with each climate variable, while still taking into account the considerable spatial variability that can be introduced from other climate conditions (e.g. a dry-warm vs. dry-cool plot) and unaccounted for environmental conditions (i.e. random effects).

### Climate and competition

Model results indicate consistent responses in both JuOs and PiEd recruitment to climate. In both species, recruitment increased, on average, as moisture availability increased, saturating or declining slightly in the wettest conditions (Figs. 1 & 2). JuOs and PiEd recruitment decreased with both increasing neighbor density and heat stress, but the overall magnitude of recruitment change due to heat stress is smaller than moisture availability and neighbor density. Both species showed consistent increases in recruitment output with plant size (Table S3 & S4).

**Figure 1.**
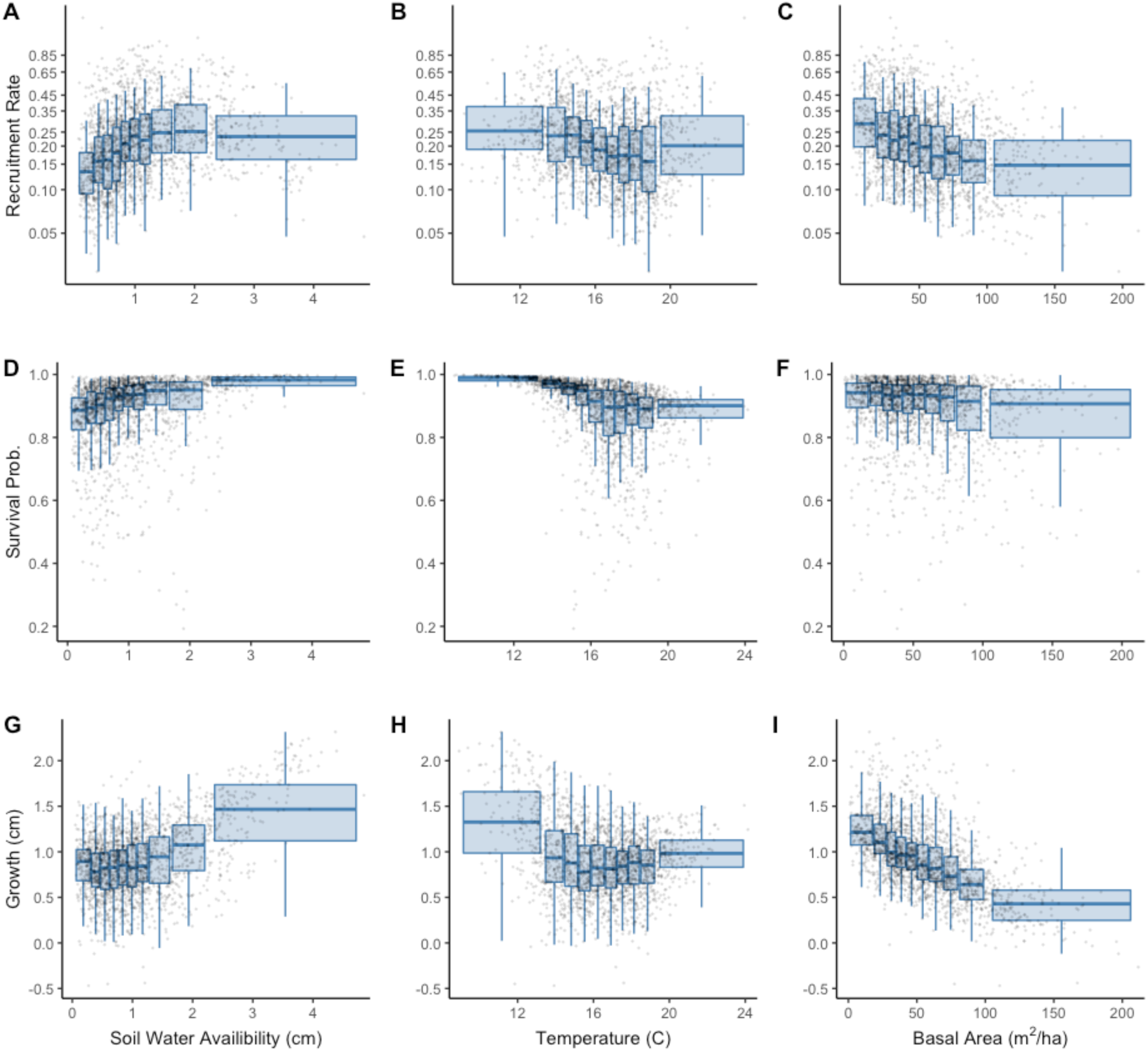
Response of *Pinus edulis* (PiEd) recruitment, survival, and size to moisture availability, heat stress, and neighbor density. Posterior mean estimates of a 15 cm diameter individual for each plot (points) are aggregated into boxplots. Growth is calculated as the change in size from the size model. Each boxplot spans a width of climate space (x-axis) that includes 10% of the total plots, i.e. each boxplot has an equal number of plots. Boxplot heights along y-axis span the spatial variability in plots created by additional climate conditions and plot random effects.

**Figure 2.**
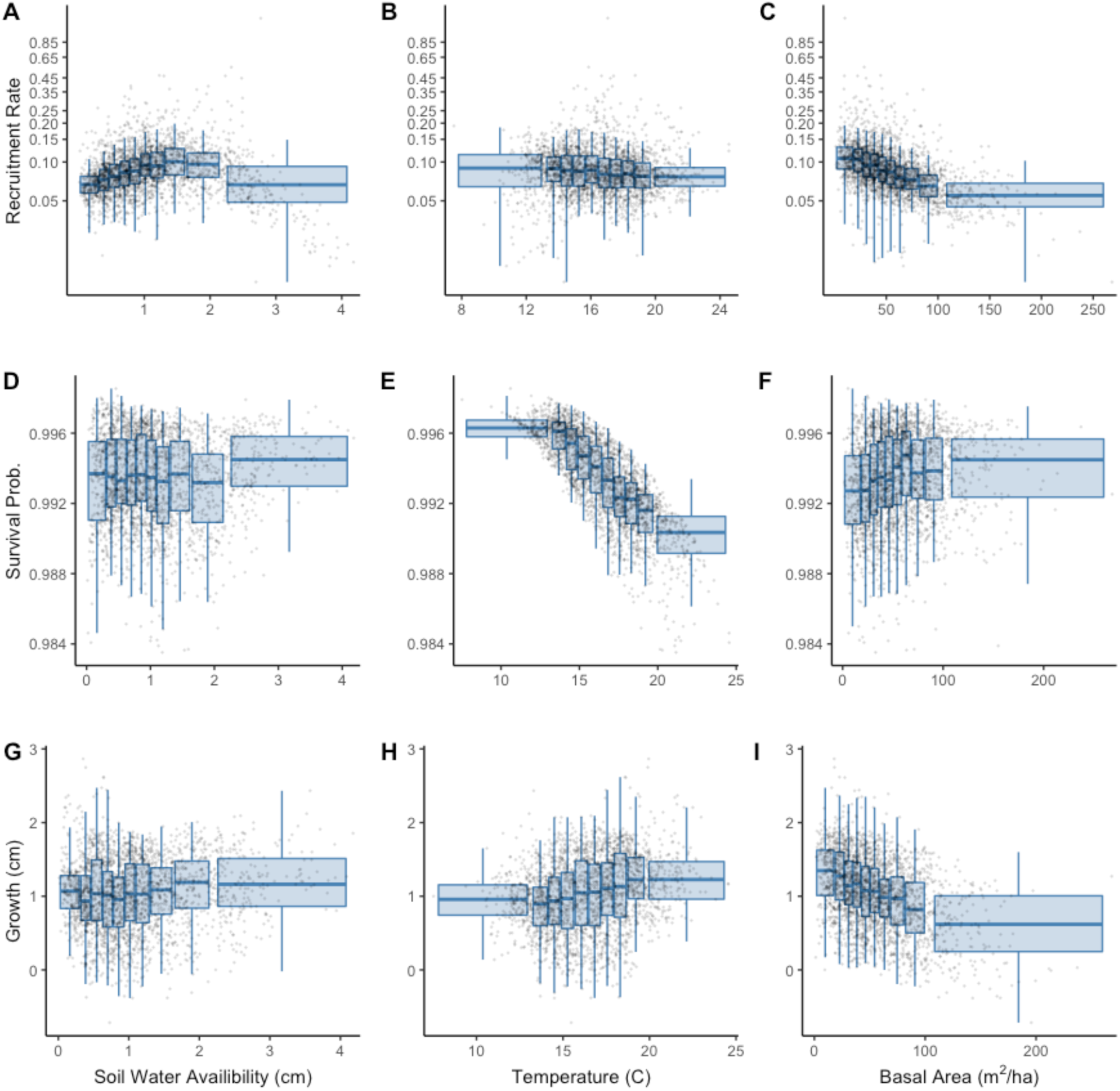
Response of *Juniperus osteosperma* (JuOs) recruitment, survival, and growth to moisture availability, heat stress, and neighbor density. Posterior mean estimates of a 15 cm diameter individual for each plot (points) are aggregated into boxplots. Growth is calculated as the change in size from the size model. Each boxplot spans a width of climate space (x-axis) that includes 10% of the total plots, i.e. each boxplot has an equal number of plots. Boxplot heights along y-axis span the spatial variability in plots created by additional climate conditions and plot random effects.

Posterior mean probabilities of survival were far more variable across plots for PiEd (0.2-1) than JuOs (0.98-1) (Figs. 1 & 2). In fact, the number of observations of JuOs mortality not associated with fire or harvest was only 153 individuals (0.6% of 25,015), compared to 1,597 (9.4% of 16,951) for PiEd. In contrast to recruitment, increasing heat stress led to clear declines in survival. JuOs and PiEd individuals in the coolest plot are expected to have 10-year survival probabilities near 1. PiEd survival rates were declined by ∼10-15% in the driest conditions, while JuOs survival declined ∼1%. Plots with higher neighbor density and lower moisture availability are also estimated to have lower PiEd survival (Fig. 3), but JuOs showed no consistent survival changes across the gradient in neighbor density or moisture availability. There was little effect of individual size on survival, and estimates overlap with 0 (Table S3 & S4).

**Figure 3.**
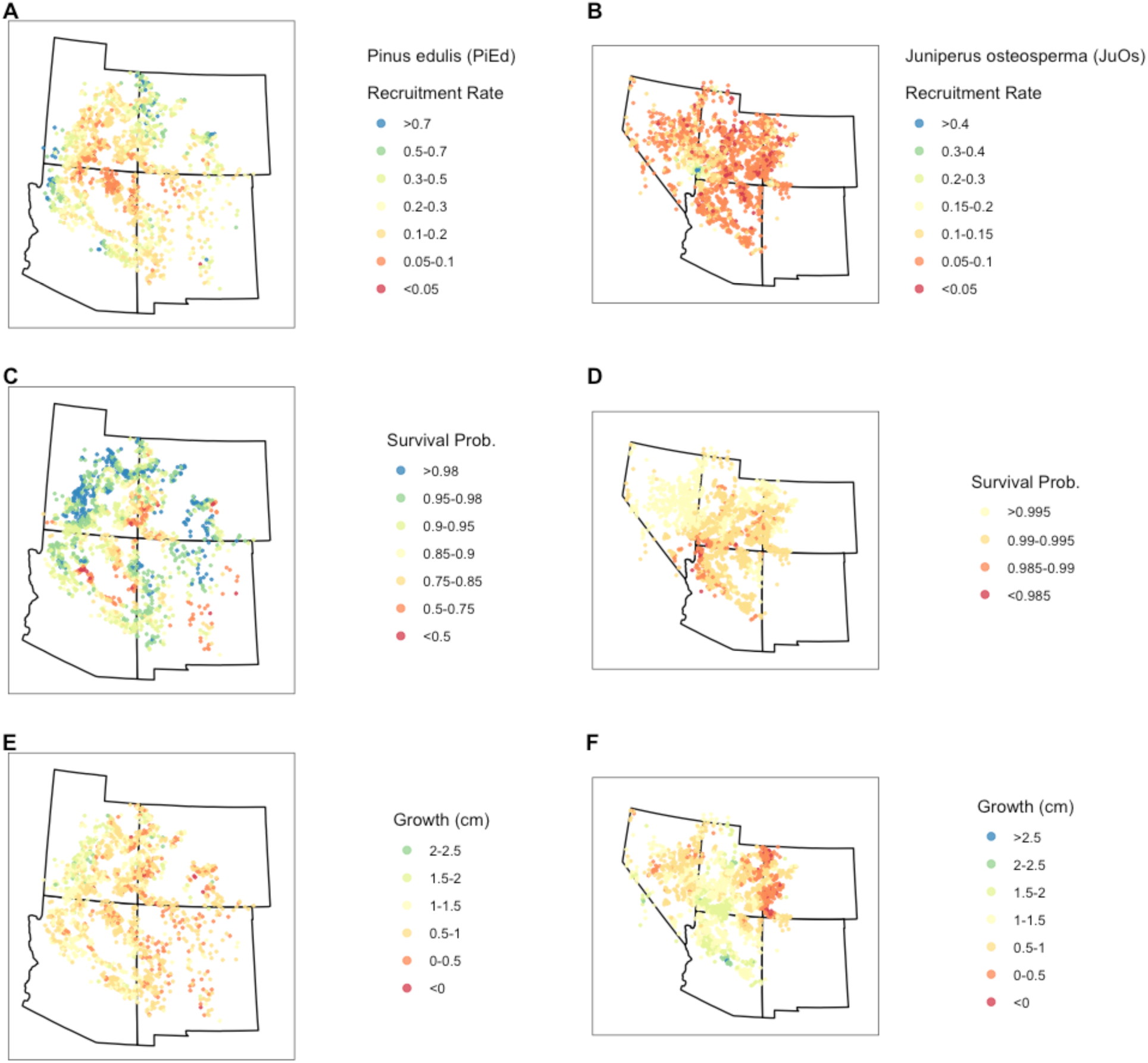
Spatial variation in recruitment, survival, and growth for PiEd and JuOs. Posterior mean estimates of a 15 cm diameter individuals recruitment rate, survival probability, and growth at all plots (points) in response to climate, competition, and unaccounted for environmental variation (random effects). Growth is calculated as the change in size from the size model. Note, points are fuzzed plot locations from publicly available FIA data.

Growth (i.e. change in diameter size) increased on average for PiEd with increasing moisture availability and declining temperatures (Fig. 1). In contrast, JuOs showed little consistent growth response to moisture availability and increasing growth with warmer temperatures (Fig. 2). Both species showed clear and consistent declines in diameter growth with increasing neighbor density and plant size, but the decrease in diameter growth with size is in part due to the radial growth geometry, and may not represent declines in overall biomass growth at larger sizes. (Table S1).

### Geographic Space

We found limited evidence of consistent responses of either species’ demographic rates to single geographic gradients (latitude, longitude, elevation), except increasing survival at high latitudes/elevations and declining growth at high elevations in JuOs (Fig. 3, S11, & S12). However, we did find clear hotspots including low survival and recruitment in PiEd in the 4 corners, lower survival and higher recruitment for JuOs in southwest Utah. We also found regions of high survival for both JuOs and PiEd in their northwestern range, with survival generally declining towards their central and southern range (Fig. 3, S11, & S12).

## Discussion

Our approach offers a promising step forward in leveraging non-traditional spatiotemporal datasets to understand the link between plant demographic rates and large scale abundance and distribution patterns (Briscoe et al. 2019). An explosion of large-scale monitoring and remote sensing dataset provide exciting opportunities to understand the drivers of species abundance and distributions, but these datasets are rarely fully compatible with traditional demographic modeling approaches (e.g. Easterling et al. 2000). For example, in our study, FIA data provide detailed information on individual growth and survival, yet recruitment of new individuals was not directly observed. To overcome this, we developed integrated modeling approach that simultaneously inferred the fate of existing seedling as well as new seedlings entering the population through recruitment. Using FIA data and novel demographic inference approaches, we found that variation in abiotic climate conditions (heat stress and moisture availability) and biotic conditions (neighbor density) can both help explain variation in the recruitment, survival, and growth of woodland species across their range. Similar modeling approaches that link individual- and population-scale data provide exciting opportunities to improve our understanding and predictions of how individual demographic rates translate into population changes over time and across landscapes (Shriver et al. 2019).

PiEd exhibited a much greater variability in survival rates, and greater sensitivity of growth and survival to heat stress and moisture availability than JuOs. This finding compliments a growing body of work which have found regional mortality in pinyon pine associated with drought and heat waves, and link the greater mortality rates in pinyon compared to juniper species to differences physiological responses to warming and drying and susceptibility of PiEd to pine beetle during drought (McDowell et al. 2008, Allen et al. 2010). Considerably less is known about environmental conditions that drive PiEd and JuOs recruitment. Consistent with findings from other semi-arid tree species (Petrie et al. 2017, Stevens-Rumann et al. 2018), we found that increasing moisture is generally associated with increasing recruitment in both JuOs and PiEd. But we also found evidence that recruitment rates may level-off, or begin to decline, in the wettest conditions. Evidence that increasing temperatures will lead directly to declining recruitment were more equivocal. With modest declines in PiEd recruitment, but no clear response in JuOs.

Together our results indicate that future warming and drying conditions, as are expected throughout the SW (Garfin et al. 2013), will likely lead to declines in survival and recruitment and increasing demographic vulnerabilities of PiEd and JuOs. Quantitatively assessing the likelihood and speed of forest decline at different locations will require integrating these demographic vulnerabilities into population models, and will be the focus of future work. We also find potential opportunities for management to alleviate the impacts of climate change. Increases in tree density lead to notable declines in all vital rates (except JuOs survival). Managed reductions in tree density could provide an opportunity to increase individual growth and decrease the risk of widespread mortality (Bradford and Bell 2017). Similarly, increases in recruitment associated with declining density could provide a natural compensatory mechanism enabling resilience of some populations following mortality events.

## Acknowledgments

This work was supported by the USGS North Central Climate Adaptation Science Center. We thank Caitlin Andrews for her assistance in preparing climate covariates. Any use of trade, firm, or product name is for descriptive purposes only and does not imply endorsement by the U.S. Government.

## Notes

https://apps.fs.usda.gov/fia/datamart/

## References

Allen, C. D., A. K. Macalady, H. Chenchouni, D. Bachelet, N. McDowell, M. Vennetier, T. Kitzberger, A. Rigling, D. D. Breshears, E. H. (Ted. Hogg, P. Gonzalez, R. Fensham, Z. Zhang, J. Castro, N. Demidova, J. H. Lim, G. Allard, S. W. Running, A. Semerci, and N. Cobb. 2010. A global overview of drought and heat-induced tree mortality reveals emerging climate change risks for forests. Forest Ecology and Management 259:660–684.

Banerjee, S., A. E. Gelfand, A. O. Finley, and H. Sang. 2008. Gaussian predictive process models for large spatial. J R Stat Soc Series B Stat Methodol. 70:825–848.

Bechtold, W. a., and P. L. Patterson. 2005. The enhanced forest inventory and analysis program – national sampling design and estimation procedures:85.

Bell, D. M., J. B. Bradford, and W. K. Lauenroth. 2014. Early indicators of change: Divergent climate envelopes between tree life stages imply range shifts in the western United States. Global Ecology and Biogeography 23:168–180.

Bradford, J. B., and D. M. Bell. 2017. A window of opportunity for climate-change adaptation: easing tree mortality by reducing forest basal area. Frontiers in Ecology and the Environment 15:11–17.

Briscoe, N. J., J. Elith, R. Salguero-Gómez, J. J. Lahoz-Monfort, J. S. Camac, K. M. Giljohann, M. H. Holden, B. A. Hradsky, M. R. Kearney, S. M. McMahon, B. L. Phillips, T. J. Regan, J. R. Rhodes, P. A. Vesk, B. A. Wintle, J. D. L. Yen, and G. Guillera-Arroita. 2019. Forecasting species range dynamics with process-explicit models: matching methods to applications. Ecology Letters 22:1940–1956.

Easterling, M. R., S. P. Ellner, P. M. Dixon, and N. Mar. 2000. Size-Specific Sensitivity : Applying a New Structured Population Model. Ecology 81:694–708.

Garfin, G., A. Jardine, R. Merideth, M. Black, and S. LeRoy. 2013. Assessment of Climate Change in the Southwest United States. National Climate Assessment.

Jackson, S. T., J. L. Betancourt, R. K. Booth, and S. T. Gray. 2009. Ecology and the ratchet of events: Climate variability, niche dimensions, and species distributions. Proceedings of the National Academy of Sciences of the United States of America 106:19685–19692.

Latimer, A. M., S. Banerjee, H. Sang, E. S. Mosher, and J. A. Silander. 2009. Hierarchical models facilitate spatial analysis of large data sets : a case study on invasive plant species in the northeastern United States. Ecology Letters 12:144–154.

McDowell, N. G., S. T. Michaletz, K. E. Bennett, K. C. Solander, C. Xu, R. M. Maxwell, and R. S. Middleton. 2018. Predicting Chronic Climate-Driven Disturbances and Their Mitigation. Trends in Ecology and Evolution 33:15–27.

McDowell, N., W. T. Pockman, C. D. Allen, D. D. Breshears, N. Cobb, T. Kolb, J. Plaut, J. Sperry, A. West, D. G. Williams, and E. a Yepez. 2008. Mechanisms of plant survival and mortality during drought: why do some plants survive while others succumb to drought? The New phytologist 178:719–39.

Millar, C. I., and N. L. Stephenson. 2015. Temperate forest health in an era of emerging megadisturbance. Science 349:823–826.

Nychka, D., and R. Furrer. 2017. fields: Tools for spatial data. R Package

Petrie, M. D., J. B. Bradford, R. M. Hubbard, W. K. Lauenroth, C. M. Andrews, and D. R. Schlaepfer. 2017. Climate change may restrict dryland forest regeneration in the 21st century. Ecology 98:1548–1559.

Rees, M., D. Z. Childs, and S. P. Ellner. 2014. Building integral projection models: A user’s guide. Journal of Animal Ecology 83:528–545.

Shriver, R. K., C. M. Andrews, R. S. Arkle, D. M. Barnard, M. C. Duniway, M. J. Germino, D. S. Pilliod, D. A. Pyke, J. L. Welty, and J. B. Bradford. 2019. Transient population dynamics impede restoration and may promote ecosystem transformation after disturbance. Ecology Letters 22: 1357–1366

Stevens-Rumann, C. S., K. B. Kemp, P. E. Higuera, B. J. Harvey, M. T. Rother, D. C. Donato, P. Morgan, and T. T. Veblen. 2018. Evidence for declining forest resilience to wildfires under climate change. Ecology Letters 21:243–252.

