## Supplemental File for "Quantifying the demographic vulnerabilities of dry woodlands to climate and competition using range-wide monitoring data"

### Supplemental Information
**Shriver et al.**

#### Covariates

Model covariates included metrics of heat stress, moisture availability, and neighbor density.

Heat stress was calculated as the average temperature (°C) during the growing season (May to October) over the 10-year period between the first and second census. Average temperatures were extracted from the Livneh gridded climate dataset (Livneh et al. 2015). The Livneh dataset provides estimates of daily temperature and precipitation at a ~7km grid cell across the United States. Because the Livneh dataset is only available through 2015, for the limited number of plots that had their second census in 2016 or 2017 the climate conditions were averaged over the 8 or 9 year period from 2006 or 2007 to 2015.

Moisture availability was calculated as the average soil water availability (SWA) in 40-100 cm depth of the soil profile during the growing season (May to October). SWA was calculated using output from SOILWAT2. SOILWAT2, a daily time-step, multiple soil layer, process-based, simulation model of ecosystem water balance that has been validated in several dryland ecosystems, (Schlaepfer et al. 2012, Bradford et al 2014), was used to model plot-specific interactions between soil conditions, vegetation, and climate. The model was run for 127,443 sites across the western US on a 1/16$^{th}$ degree grid. This spatial resolution and grid is the same as the underlying climate data (Livneh et al. 2015) that drove the SOILWAT2 simulations. Daily climate data, including daily maximum and minimum temperatures and precipitation was obtained for 1915–2015.

Soils data for the model was extracted from the ISRIC Wise Global v1.2 dataset (Batjes 2012) and consisted of texture (%clay and % sand), gravel (% volume), and matric density (g/cm$^3$) for 8 soil layers (10, 20, 40, 60, 80, 100, 150, 200 cm), or until maximum depth was reached, at each site. Vegetation composition in SOILWAT2 was parameterized, including static estimates of biomass and phenology for different plant function types (PFTs), were generated using algorithms that account for the effect of temperature and precipitation on these factors under both current and future climate scenarios (see Paruelo and Lauenroth 1996, Bradford et al. 2014 for more details).

These data were formatted and ingested into the SOILWAT2 ecohydrological model using the rSOILWAT2 (Schlaepfer and Murphy 2018) and rSFSW2 (Schlaepfer and Andrews. 2018) R (R Core Team, 2019) packages. From the outputs of the SOILWAT2 model, average annual soil water availability (SWA, cm) was calculated in R. SWA was calculated by summing SWA across every layer per site, where the soil water available per layer was calculated as the difference between actual volumetric water content (VWC, cm/cm) and the amount of VWC potentially available at a soil water potential threshold of -3.9 MPa.

Neighbor density was calculated as the basal area (m$^2$/hectare) in each plot. Basal area data was extracted from the FIA dataset from all living trees (regardless of species) in the first census and correcting for differing sampling plot sizes.

Covariates were centered and standardized by mean and variance to improve model convergence. Models were fit using the original, raw data scale (inches) for plant size and results were converted to metric units.

**Further modeling details**

Full posterior distribution for each species is

$$[\mathbf{b}_{(z)}, \mathbf{b}_{(s)}, \mathbf{b}_{(f)}, \sigma, \nu, \upsilon, \boldsymbol{\delta}, \boldsymbol{\omega}, \boldsymbol{\gamma} | \mathbf{z}, \mathbf{s}, \mathbf{c}'] = \prod_i [\mathrm{z}_{\boldsymbol{i},\boldsymbol{t}+\mathbf{1}} | g(\alpha_{(z)}, z_{i,t}, \mathbf{X}_{d[i]}, \mathbf{b}_{(z)}, \omega_{d[i]}), \sigma]$$

$$\prod_i [\mathrm{s}_{\boldsymbol{i},\boldsymbol{t}+\mathbf{1}} | g(\alpha_{(s)}, z_{i,t}, \mathbf{X}_{d[i]}, \mathbf{b}_{(s)}, \delta_{d[i]})]$$

$$\prod_d [\mathbf{c}'_{d,t+1} | g(\alpha_{(s)}, \mathbf{b}_{(s)}, \alpha_{(z)}, \mathbf{b}_{(z)}, \alpha_{(f)}, \mathbf{b}_{(f)}, \alpha_{(f)}, \mathbf{X}_d, m_j, \sigma, \nu, \upsilon, \delta_d, \omega_d, \gamma_{\boldsymbol{d}}, \mathbf{n}_{d,t}), \kappa]$$

$$[\boldsymbol{\delta} | \tau_{(s)}, \phi_{(s)}][\boldsymbol{\omega} | \tau_{(z)}, \phi_{(z)}][\boldsymbol{\gamma} | \tau_{(f)}, \phi_{(f)}]$$

$$[\alpha_{(s)}][\alpha_{(z)}][\alpha_{(f)}][\mathbf{b}_{(s)}][\mathbf{b}_{(z)}][\mathbf{b}_{(f)}][\sigma][\nu][\upsilon][\tau_{(s)}][\tau_{(z)}][\tau_{(f)}][\phi_{(s)}][\phi_{(z)}][\phi_{(f)}][\kappa]$$

**Computation**

We fit our models with Hamiltonian Monte Carlo (HMC) using the "rstan" package (Stan Development Team 2020). Models were run with 3 chains and 2000 iterations each, 1000 of which were warmup. HMC algorithms are more efficient at exploring posterior parameter space, per iteration, than traditional MCMC algorithms, and thus require substantially fewer iterations to reach convergence. Parameter convergence was monitored visually and using convergence statistics (R-hat) and model fits were also evaluated with posterior predictive checks and posterior p-values (See below, Figs. S5-S10)

**Random effect details**

*Knot locations*

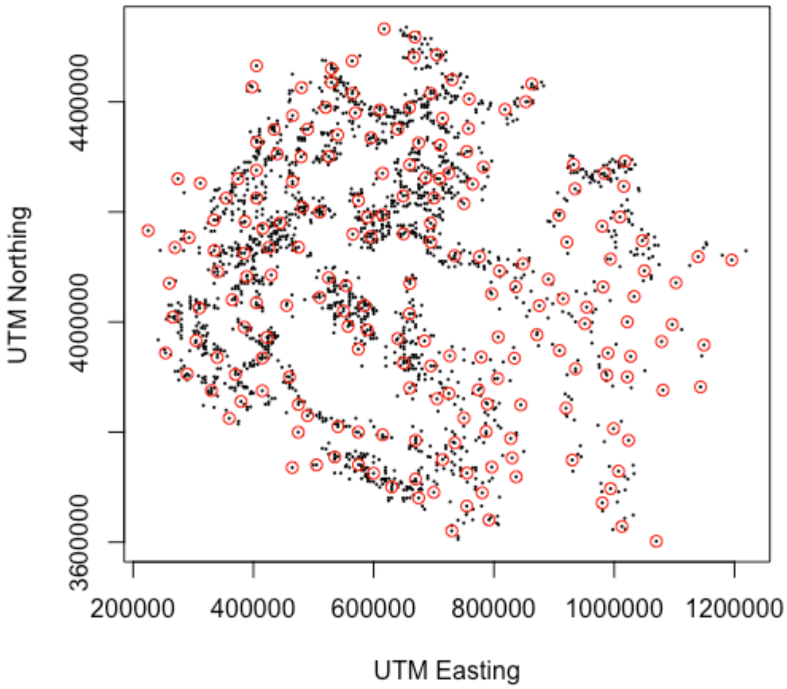


**Figure S1. Locations of *Pinus edulis* (PiEd) knots (red points) and fuzzed locations of plots based on publicly available FIA data (black points).** Knot locations were determined to maximize spatial coverages using the “fields” package in R.

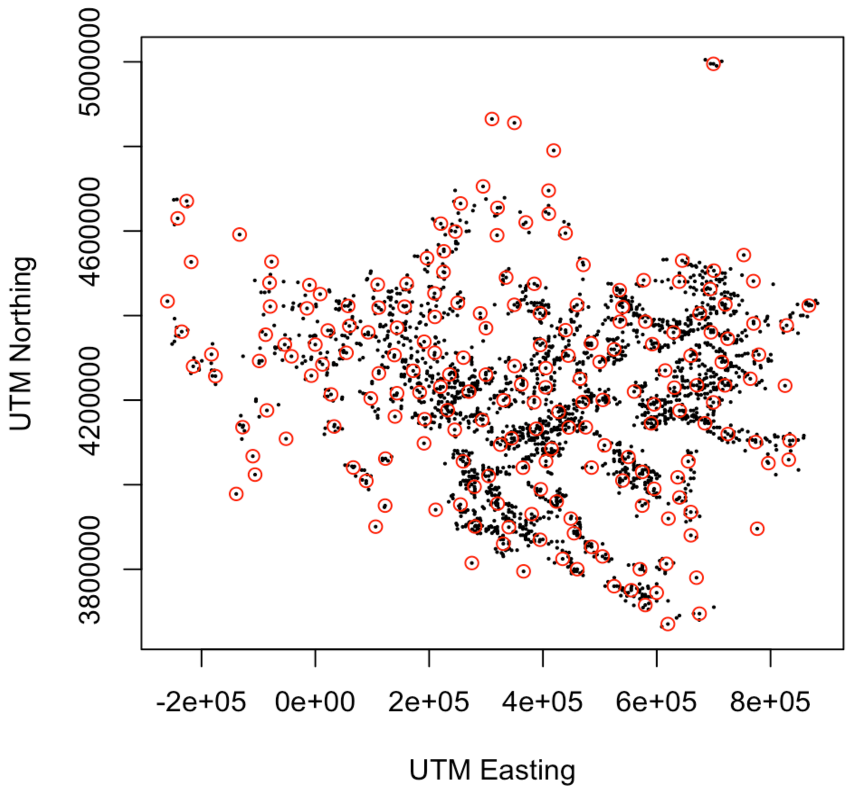


**Figure S2. Locations of *Juniperus osteosperma* (JuOs) knots (red points) and fuzzed locations of plots based on publicly available FIA data (black points).** Knot locations were determined to maximize spatial coverages using the “fields” package in R.

*Random effect estimates*

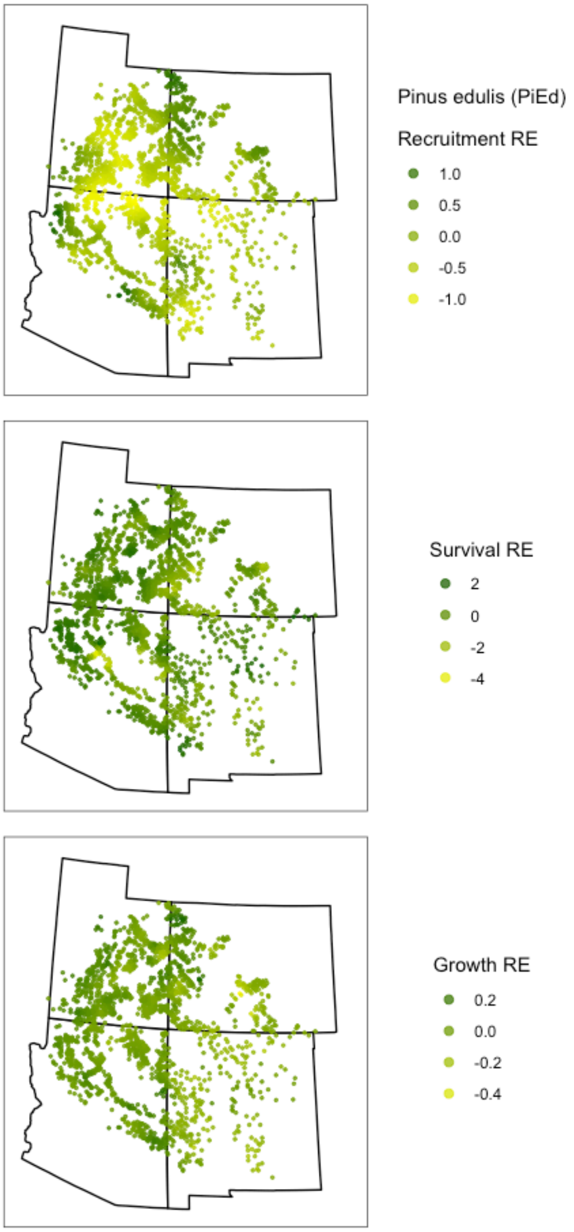


**Figure S3. Posterior mean random effect values for PiEd.** Values are on the linear scale, untransformed by link functions. Note, points are fuzzed plot locations from publicly available FIA data.

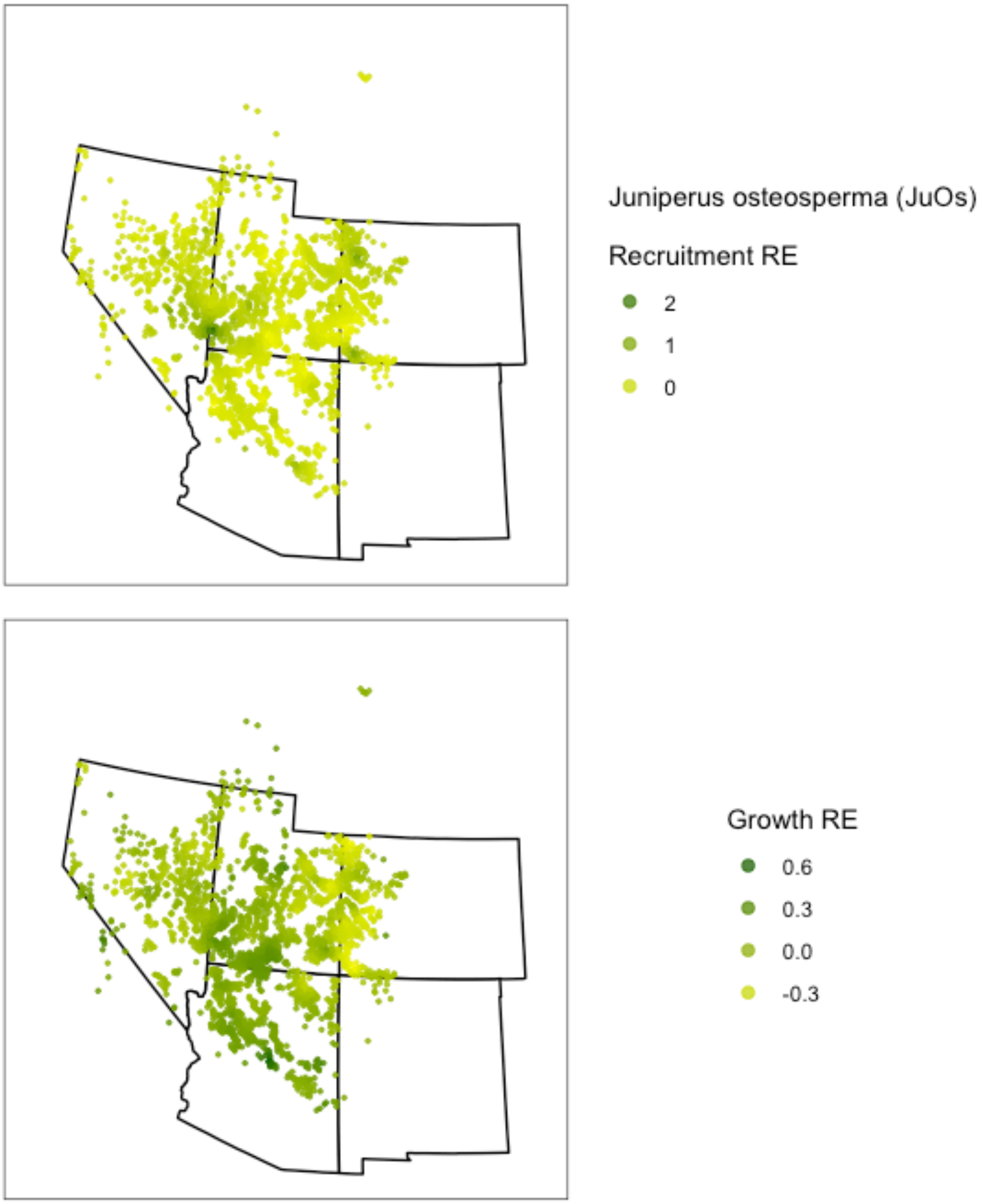


**Figure S4. Posterior mean random effect values for JuOs.** Values are on the linear scale, untransformed by link functions. Note, points are fuzzed plot locations from publicly available FIA data.

**Posterior Checks**

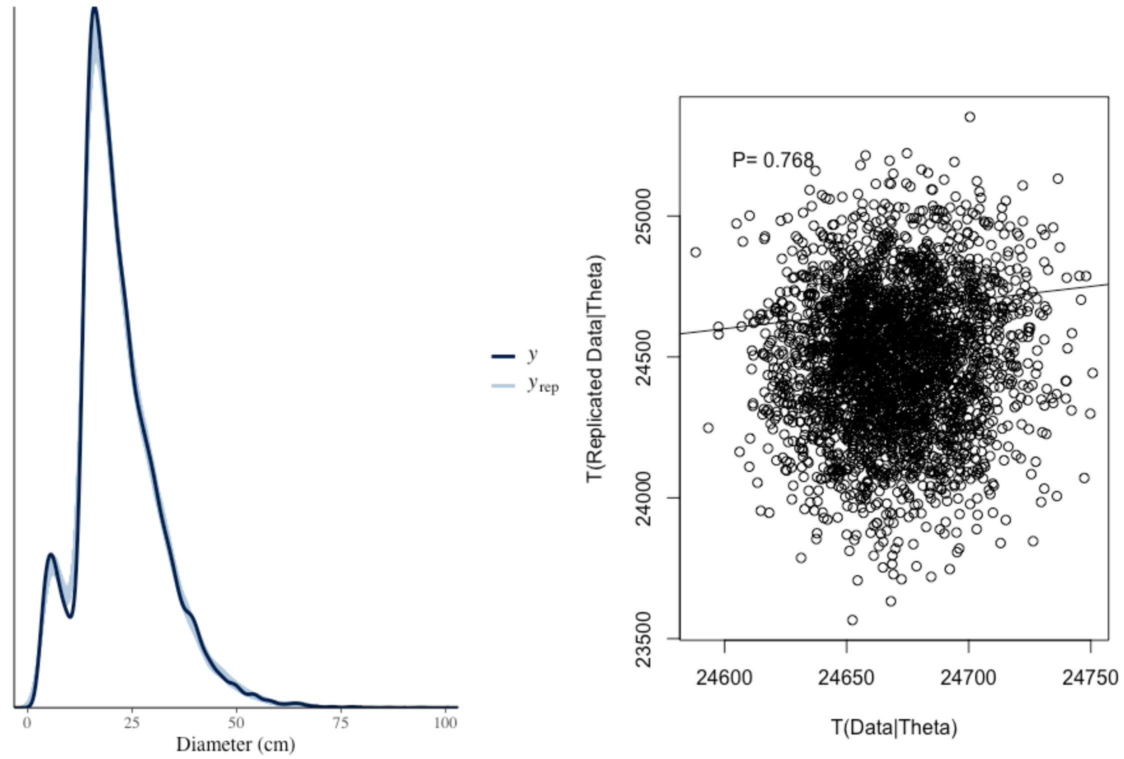


**Figure S5. PiEd posterior predictive checks for plant size at *t+1*.** Posterior predictive checks indicate good agreement between model predictions ($y_{rep}$) and observed data (y). Posterior predictive p-values assess model fit by comparing replicated data from the fit model to the real measured data using a test function. P-values <0.05 or >0.95 indicate high probability that model predictions are more extreme than real data, and thus poorer model fit (Gelman et al. 2004). We use a deviance test function for p-values.

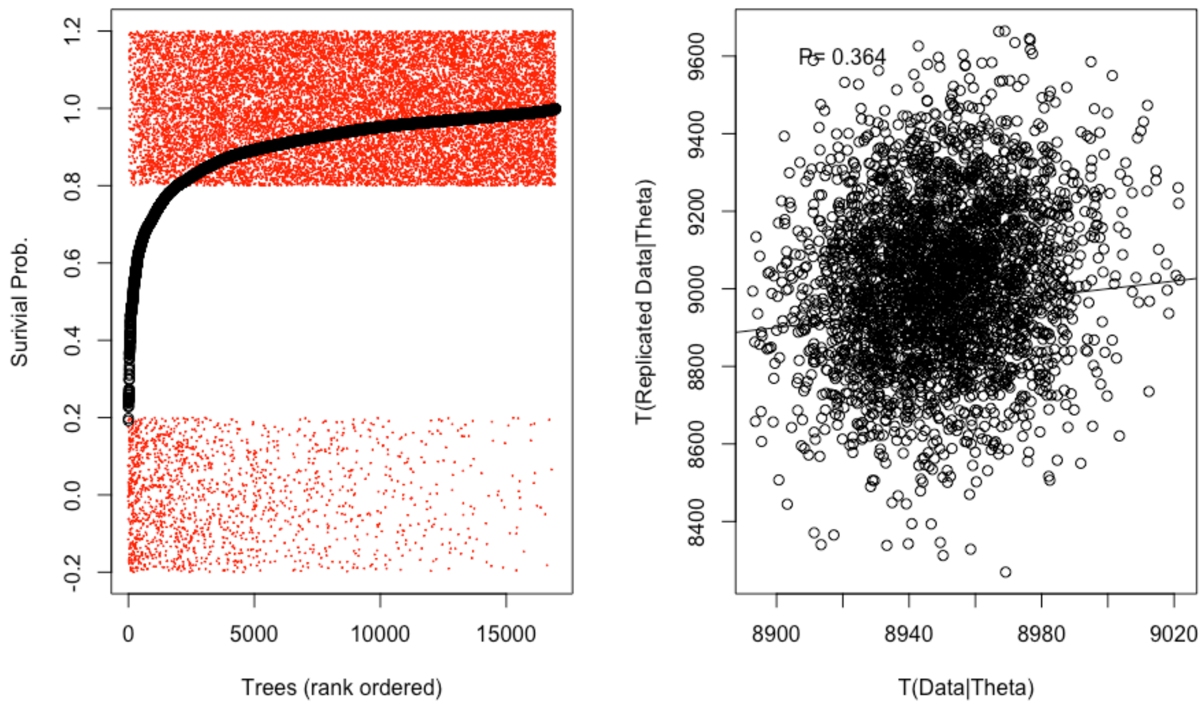


**Figure S6. PiEd posterior predictive checks for plant survival.** Posterior predictive checks indicate good agreement between mean model predictions (black points) and observed data (jittered red points, 1=Alive, 0=Dead). Note: 95% CI are omitted and red points are 'jittered' for clarity. Posterior predictive p-values assess model fit by comparing replicated data from the fit model to the real measured data using a test function. P-values <0.05 or >0.95 indicate high probability that model predictions are more extreme than real data, and thus poorer model fit (Gelman et al. 2004). We use a deviance test function for p-values.

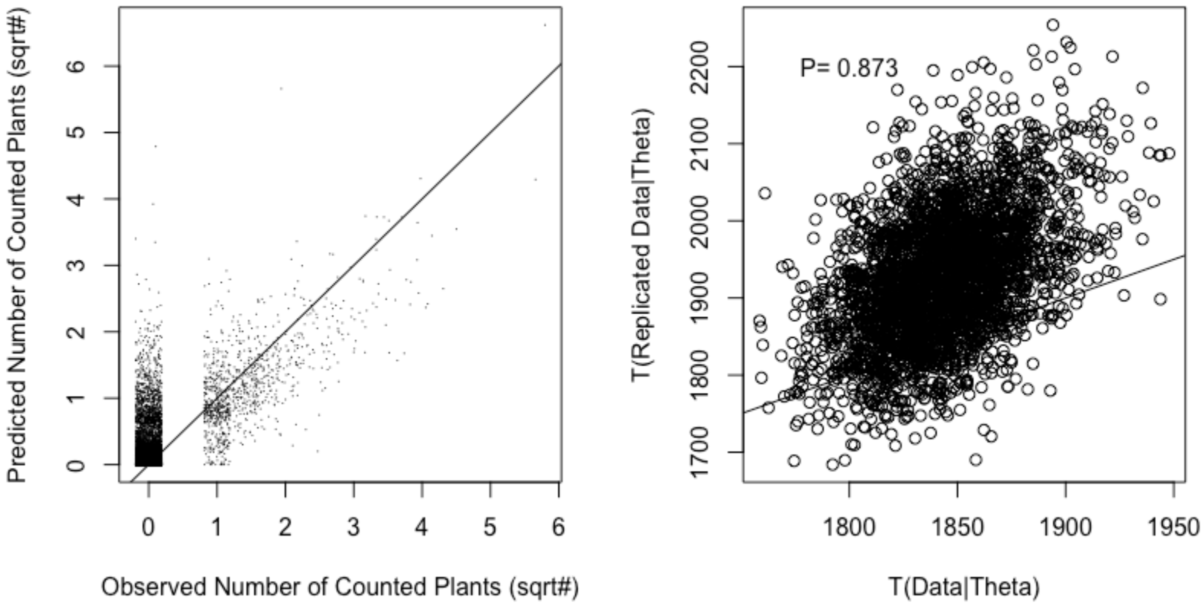


**Figure S7. PiEd posterior predictive checks for plant recruitment.** Posterior predictive checks indicate good agreement between mean model predictions (y-axis, left panel) and observed data (x-axis, left panel). Note: 95% CI are omitted for clarity. Posterior predictive p-values assess model fit by comparing replicated data from the fit model to the real measured data using a test function. P-values <0.05 or >0.95 indicate high probability that model predictions are more extreme than real data, and thus poorer model fit (Gelman et al. 2004). We use a Freeman-Tukey test function, intended for count data with low expected values, for p-values.

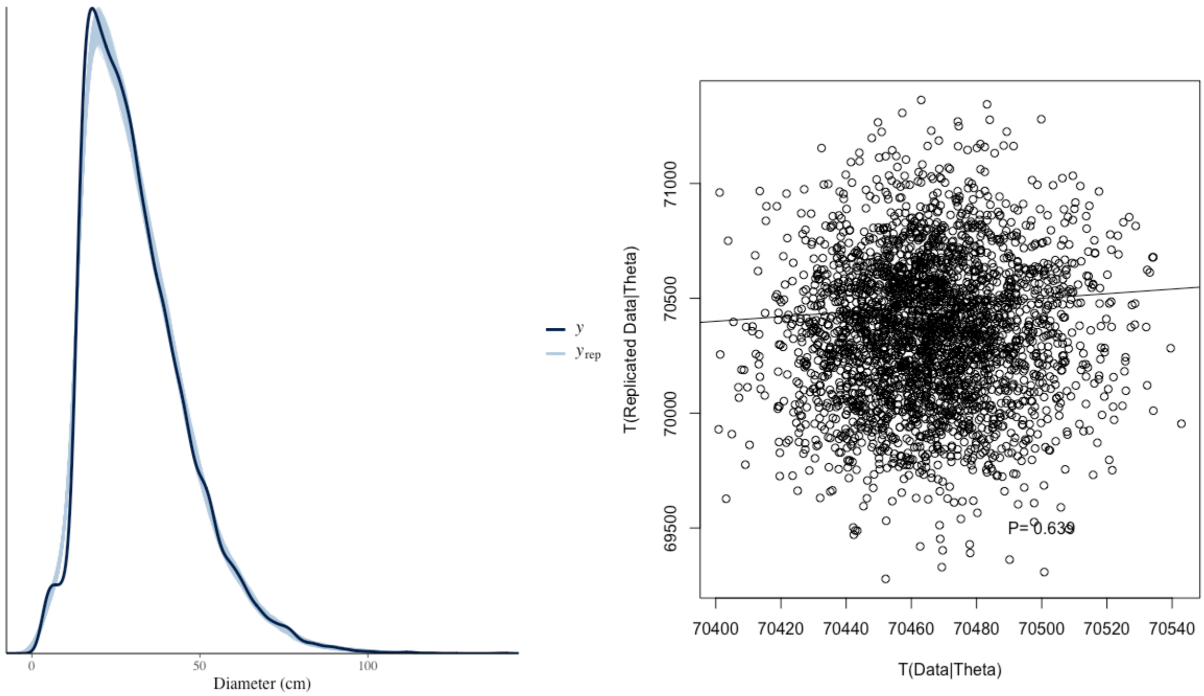


**Figure S8. JuOs posterior predictive checks for plant size at *t+1*.** Posterior predictive checks indicate good agreement between model predictions ($y_{rep}$) and observed data (y). Posterior predictive p-values assess model fit by comparing replicated data from the fit model to the real measured data using a test function. P-values <0.05 or >0.95 indicate high probability that model predictions are more extreme than real data, and thus poorer model fit (Gelman et al. 2004). We use a deviance test function for p-values.

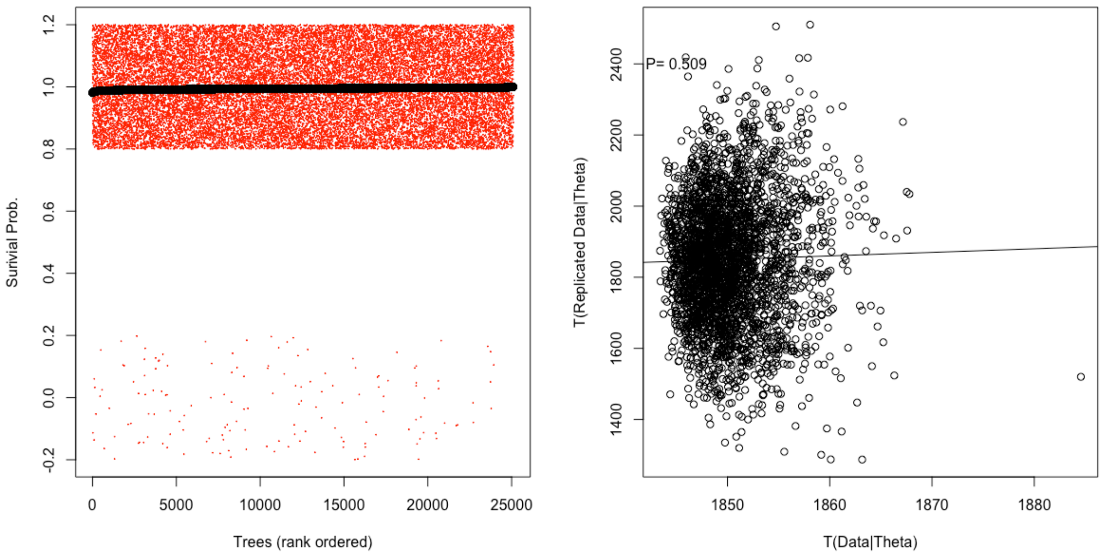


**Figure S9. JuOs posterior predictive checks for plant survival.** Posterior predictive checks indicate good agreement between mean model predictions (black points) and observed data (jittered red points, 1=Alive, 0=Dead). Note: 95% CI are omitted and red points are ‘jittered’ for clarity. Posterior predictive p-values assess model fit by comparing replicated data from the fit model to the real measured data using a test function. P-values <0.05 or >0.95 indicate high probability that model predictions are more extreme than real data, and thus poorer model fit (Gelman et al. 2004). We use a deviance test function for p-values.

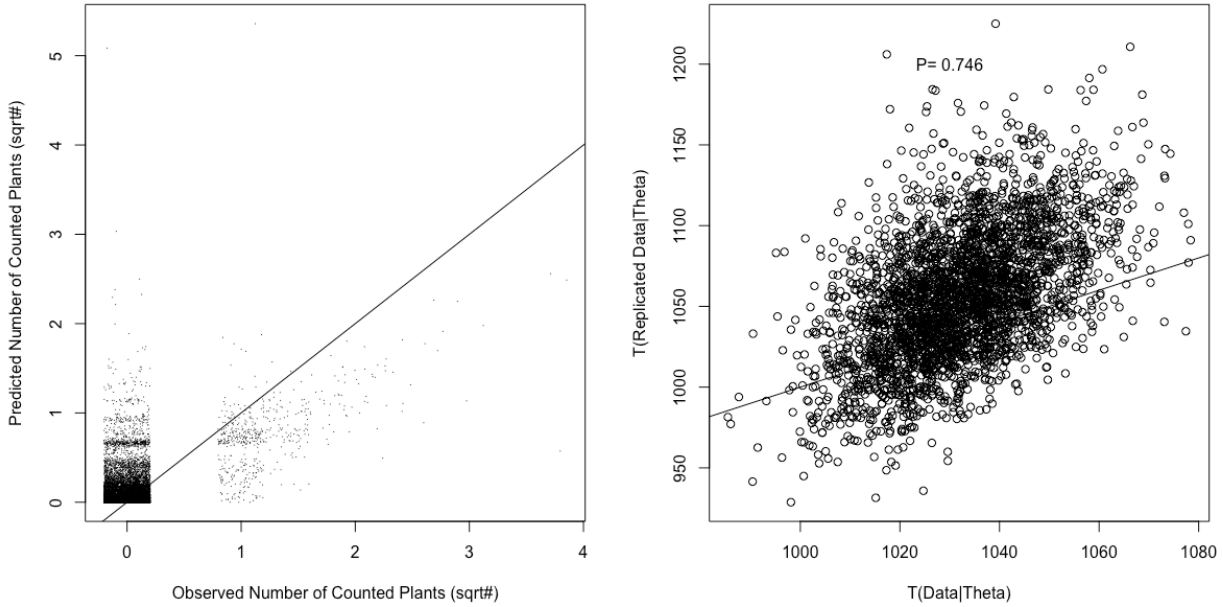


**Figure S10. JuOs posterior predictive checks for plant recruitment.** Posterior predictive checks indicate good agreement between mean model predictions (y-axis, left panel) and observed data (x-axis, left panel). Note: 95% CI are omitted for clarity. Posterior predictive p-values assess model fit by comparing replicated data from the fit model to the real measured data using a test function. P-values <0.05 or >0.95 indicate high probability that model predictions are more extreme than real data, and thus poorer model fit (Gelman et al. 2004). We use a Freeman-Tukey test function, intended for count data with low expected values, for p-values.

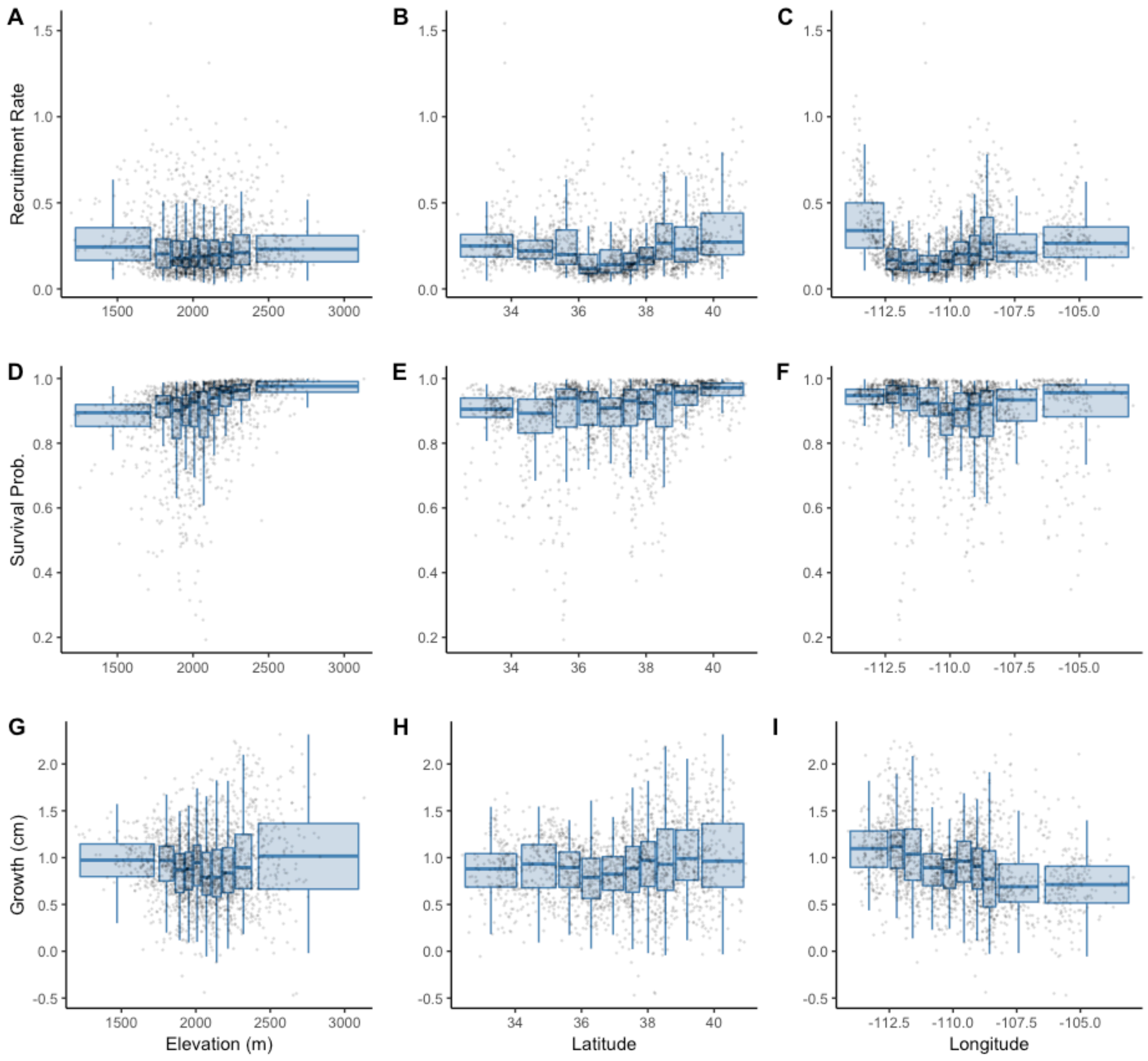


**Figure S11. Response of *Pinus edulis* (PiEd) recruitment, survival, and size to elevation, latitude, and longitude.** Posterior mean estimates of a 15 cm diameter individual for each plot (points) are aggregated into boxplots. Each boxplot spans a width of space (x-axis) that includes 10% of the total plots, i.e. each boxplot has an equal number of plots. Boxplot heights along y-axis span the spatial variability in plots created by climate conditions and plot random effects.

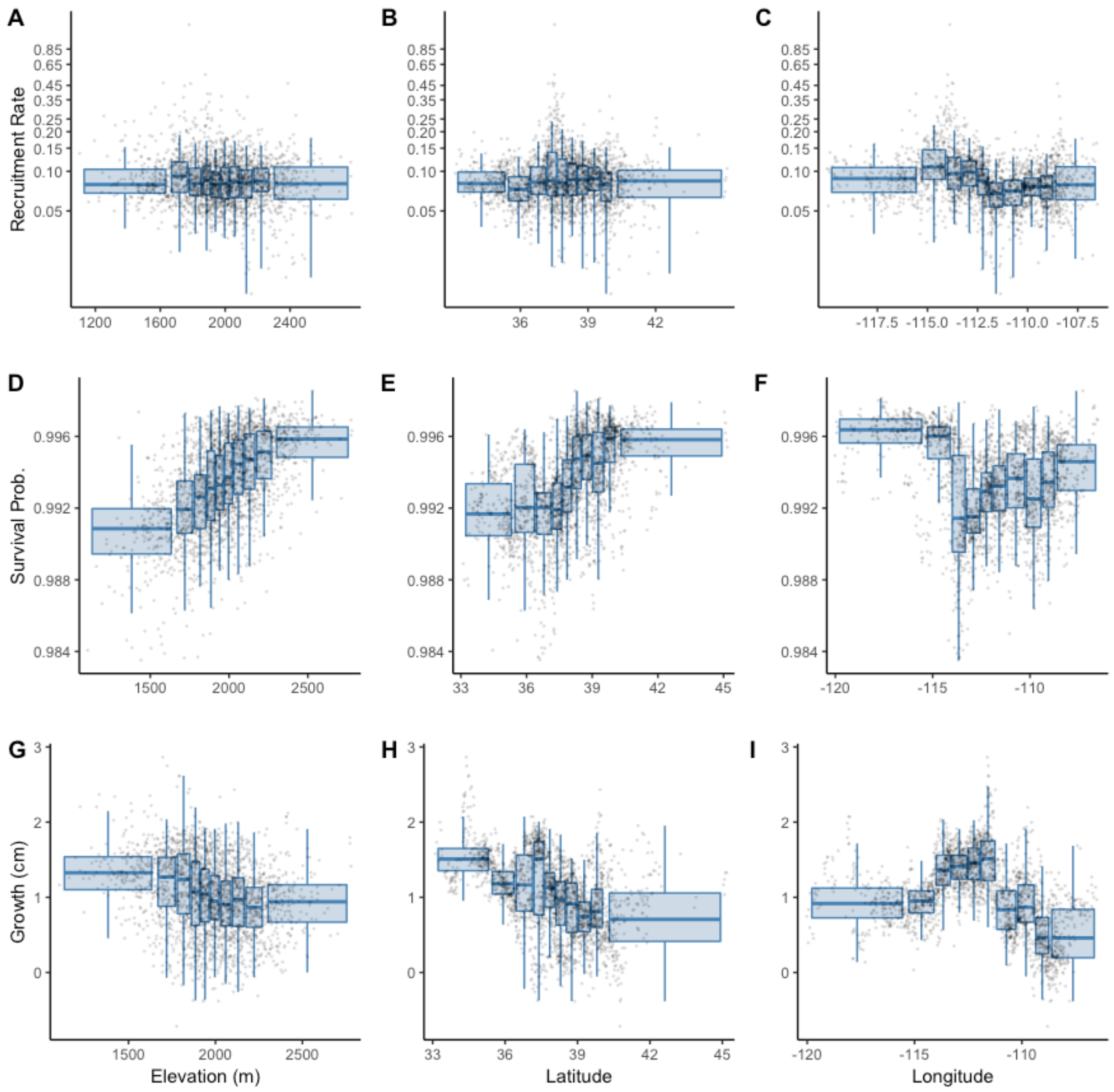


**Figure S12. Response of *Juniperus osteosperma* (JuOs) recruitment, survival, and size to elevation, latitude, and longitude.** Posterior mean estimates of a 15 cm diameter individual for each plot (points) are aggregated into boxplots. Each boxplot spans a width of space (x-axis) that includes 10% of the total plots, i.e. each boxplot has an equal number of plots. Boxplot heights along y-axis span the spatial variability in plots created by climate conditions and plot random effects.

**Priors**

**Table S1. PiEd model priors**

| Parameter | Distribution | Values |
|---|---|---|
| $\mathbf{b}_{(s)}, \mathbf{b}_{(z)}, \mathbf{b}_{(f)}, \nu$ | Normal | mu=0, sigma=5 |
| $\tau_{(s)}, \tau_{(z)}, \tau_{(f)}$ | Half-Normal | mu=0, sigma =5 |
| $\phi_{(s)}, \phi_{(z)}, \phi_{(f)}$ | Half-Normal | mu=0, sigma =5 |
| $\upsilon$ | Half-Cauchy | x_0=0, gamma=1 |
| $\kappa$ | Half-Cauchy | x_0=0, gamma=5 |
| $\sigma$ | Half-Cauchy | x_0=0, gamma=5 |

**Table S2. JuOs model priors**

| Parameter | Distribution | Values |
|---|---|---|
| $\mathbf{b}_{(s)\ [excpt.Density]}, \mathbf{b}_{(z)}, \mathbf{b}_{(f)}, \nu$ | Normal | mu=0, sigma=5 |
| $\mathbf{b}_{(s)\ [Density]}$ | Normal | mu=0, sigma=1 |
| $\tau_{(z)}, \tau_{(f)}$ | Half-Normal | mu=0, sigma =5 |
| $\phi_{(z)}, \phi_{(f)}$ | Half-Normal | mu=0, sigma =5 |
| $\upsilon$ | Half-Cauchy | x_0=0, gamma=1 |
| $\kappa$ | Half-Cauchy | x_0=0, gamma=5 |
| $\sigma$ | Half-Cauchy | x_0=0, gamma=5 |

#### Parameter Estimates

**Table S3. PiEd parameter estimates**

| Parameter | Mean | 2.50% | 97.50% |
|---|---|---|---|
| $\mathbf{b}_{(s)}$-Int. | 2.58 | 2 | 3.318 |
| $\mathbf{b}_{(s)}$-Size | -0.021 | -0.036 | -0.006 |
| $\mathbf{b}_{(s)}$-Temp. | -0.911 | -1.102 | -0.725 |
| $\mathbf{b}_{(s)}$-Precip. | 0.058 | -0.103 | 0.221 |
| $\mathbf{b}_{(s)}$-Temp$^2$. | 0.176 | 0.082 | 0.274 |
| $\mathbf{b}_{(s)}$-Precip$^2$. | -0.062 | -0.135 | 0.014 |
| $\mathbf{b}_{(s)}$-Dense. | -0.279 | -0.343 | -0.217 |
| $\mathbf{b}_{(z)}$-Int. | 0.435 | 0.39 | 0.477 |
| $\mathbf{b}_{(z)}$-Size | 0.979 | 0.977 | 0.981 |
| $\mathbf{b}_{(z)}$-Temp. | -0.025 | -0.047 | -0.002 |
| $\mathbf{b}_{(z)}$-Precip. | 0.035 | 0.013 | 0.057 |
| $\mathbf{b}_{(z)}$-Temp$^2$. | 0.018 | 0.007 | 0.028 |
| $\mathbf{b}_{(z)}$-Precip$^2$. | 0.011 | 0.003 | 0.02 |
| $\mathbf{b}_{(z)}$-Dense. | -0.095 | -0.105 | -0.085 |
| $\sigma$ | 0.537 | 0.531 | 0.543 |
| $\mathbf{b}_{(f)}$-Int. | -1.892 | -2.581 | -0.992 |
| $\mathbf{b}_{(f)}$-Size | 0.089 | 0.058 | 0.118 |
| $\mathbf{b}_{(f)}$-Temp. | 0.013 | -0.206 | 0.231 |
| $\mathbf{b}_{(f)}$-Precip. | 0.364 | 0.118 | 0.62 |
| $\mathbf{b}_{(f)}$-Temp$^2$. | 0.063 | -0.056 | 0.174 |
| $\mathbf{b}_{(f)}$-Precip$^2$. | -0.176 | -0.297 | -0.063 |
| $\mathbf{b}_{(f)}$-Dense. | -0.268 | -0.397 | -0.143 |
| $\phi_{(z)}$ | 2.79 | 1.557 | 4.129 |
| $\phi_{(f)}$ | 1.056 | 0.118 | 2.378 |
| $\phi_{(s)}$ | 1.787 | 1.021 | 2.527 |
| $\tau_{(z)}$ | 0.027 | 0.018 | 0.038 |
| $\tau_{(f)}$ | 1.002 | 0.384 | 3.048 |
| $\tau_{(s)}$ | 2.703 | 1.848 | 4.083 |
| $\upsilon$ | 2.762 | 2.268 | 3.279 |
| $\nu$ | -14.582 | -20.717 | -9.512 |
| $\kappa$ | 0.932 | 0.792 | 1.096 |

**Table S4. JuOs parameter estimates**

| Parameter | Mean | 2.50% | 97.50% |
|---|---|---|---|
| $\mathbf{b}_{(s)}$-Int. | 4.966 | 4.557 | 5.358 |
| $\mathbf{b}_{(s)}$-Size | 0.01 | -0.017 | 0.04 |
| $\mathbf{b}_{(s)}$-Temp. | -0.502 | -0.733 | -0.279 |
| $\mathbf{b}_{(s)}$-Precip. | -0.355 | -0.614 | -0.101 |
| $\mathbf{b}_{(s)}$-Temp$^2$. | 0.026 | -0.123 | 0.195 |
| $\mathbf{b}_{(s)}$-Precip$^2$. | 0.071 | -0.066 | 0.221 |
| $\mathbf{b}_{(s)}$-Dense. | 0.001 | -0.144 | 0.154 |
| $\mathbf{b}_{(z)}$-Int. | 0.663 | 0.583 | 0.754 |
| $\mathbf{b}_{(z)}$-Size | 0.963 | 0.961 | 0.965 |
| $\mathbf{b}_{(z)}$-Temp. | 0.006 | -0.031 | 0.041 |
| $\mathbf{b}_{(z)}$-Precip. | 0.024 | -0.005 | 0.053 |
| $\mathbf{b}_{(z)}$-Temp$^2$. | -0.012 | -0.028 | 0.003 |
| $\mathbf{b}_{(z)}$-Precip$^2$. | 0.002 | -0.01 | 0.014 |
| $\mathbf{b}_{(z)}$-Dense. | -0.049 | -0.061 | -0.036 |
| $\sigma$ | 0.991 | 0.983 | 1 |
| $\mathbf{b}_{(f)}$-Int. | -2.685 | -3.216 | -2.22 |
| $\mathbf{b}_{(f)}$-Size | 0.036 | 0.001 | 0.07 |
| $\mathbf{b}_{(f)}$-Temp. | 0.003 | -0.237 | 0.256 |
| $\mathbf{b}_{(f)}$-Precip. | 0.175 | -0.098 | 0.438 |
| $\mathbf{b}_{(f)}$-Temp$^2$. | 0.027 | -0.133 | 0.176 |
| $\mathbf{b}_{(f)}$-Precip$^2$. | -0.184 | -0.361 | -0.023 |
| $\mathbf{b}_{(f)}$-Dense. | -0.239 | -0.406 | -0.078 |
| $\phi_{(z)}$ | 1.596 | 0.764 | 2.556 |
| $\phi_{(f)}$ | 3.803 | 1.048 | 6.642 |
| $\tau_{(z)}$ | 0.057 | 0.039 | 0.083 |
| $\tau_{(f)}$ | 0.837 | 0.292 | 1.734 |
| $\upsilon$ | 2.75 | 2.137 | 3.412 |
| $\nu$ | -11.249 | -17.642 | -6.276 |
| $\kappa$ | 0.73 | 0.572 | 0.926 |

**References**


Batjes, N. H. 2012. ISRIC-WISE derived soil properties on a 5 by 5 arc-minutes global grid (ver. 1.2). ISRIC-World Soil Information.

Bradford, J. B., D. R. Schlaepfer, and W. K. Lauenroth. 2014. Ecohydrology of Adjacent Sagebrush and Lodgepole Pine Ecosystems: The Consequences of Climate Change and Disturbance. Ecosystems.

Gelman, A., J. Carlin, H. Stern, and D. Rubin. 2004. Bayesian Data Analysis. Chapman and Hall/CRC.

Livneh, B., T. J. Bohn, D. W. Pierce, F. Munoz-Arriola, B. Nijssen, R. Vose, D. R. Cayan, and L. Brekke. 2015. A spatially comprehensive, hydrometeorological data set for Mexico, the U.S., and Southern Canada 1950-2013. Scientific Data 2:1–12.

Paruelo, J., and W. Lauenroth. 1996. Relative abundance of plant functional types in grasslands and shrublands of north America. Ecological Applications 6:1212–1224.

Schlaepfer, D. R., and C. A. Andrews. 2018. rSFSW2: Simulation Framework for SOILWAT2 R package.

Schlaepfer, D. R., and R. Murphy. 2018. rSOILWAT2: An Ecohydrological Ecosystem-Scale Water Balance Simulation Model R package .

Stan Development Team. 2020. RStan: the R interface to Stan.
